# Rice novel useful semidwarf gene *d60* on chromosome 2 causing pleiotropically gamete abortion

**DOI:** 10.1101/2024.06.26.600879

**Authors:** Motonori Tomita, Daisuke Kamiya, Keisuke Okawa, Kohei Nakayama

## Abstract

Rice semidwarf gene *d60* is inherited according to a unique mode because it causes pleiotropically gamete abortion in the case of coexistence with the gamete lethal gene *gal*. Namely, F_2_ progenies of Koshihikari (*D60D60galgal*, long stem) × Hokuriku 100 (*d60d60GalGal*, short stem) segregated in the ratio of 1 semidwarf (1 *d60d60GalGal*):2 tall and quarter sterile (2 *D60d60Galgal*):6 tall (2 *D60d60GalGal*:1 *D60D60GalGal*:2 *D60D60Galgal*:1 *D60D60galgal*), which is skewed from the Mendelian 1 semidwarf:3 tall ratio. Firstly, genetic linkage was tested on the basis that the segregation ratios of the phenotypic marker genes linked to *d60* do not fit Mendelian ratio. F_2_ of Koshihikari d60 line (*d60d60GalGal)* × lines carrying *d30* or *gh2* on chromosome 2, the segregation ratios of these alleles were deviated from 1:3 but well fitted to 5:4 when *d30 and gh2* were closely linked to *D60*. We then conducted molecular fine mapping of *d60* by using DNA polymorphisms in F_2_ of Koshihikari d60 line × line carrying the indica chromosome 2 segment in Koshihikari. As a result, *d60* was tightly linked with RM12970 by a recombination value 0.0 at the region 10.3 Mb away from the distal end of the short arm of chromosome 2. Whole genome sequencing of Koshihikari d60 revealed no SNPs and Indels around the 10.3 Mb region. RT-qPCR for ent-copalyl diphosphate synthase-like gene on 10.3 Mb indicated its transcription was reduced compared to that of Koshihikari.

---

Rice (*Oryza sativa* L.) is one of the most important food crops, feeding half of the world’s population (Fairhurst and Dobermann 2002), particularly those living in the monsoon areas of Asia. The development of high-yielding semidwarf varieties of rice led to a rapid increase in the global production of cereal grains, which more than doubled from 1960 to 1990 (Khush 1990). Widespread adoption of miracle rice released by the International Rice Research Institute (IRRI) brought about a globally “green revolution”, particularly in the monsoonal regions of Asia, where typhoons frequently occur during the harvesting season (Khush 1990; Athwal 1971). Dwarfing prevents plants from lodging at their full-ripe stage, makes them lodging-resistant to wind and rain, has enhanced their adaptability for heavy manuring, and has dramatically improved the productivity of rice. Semidwarf gene *sd1* contributed to the ‘Green Revolution’ of rice, encodes a defective Gibbelleric Acid (GA) 20-oxidase gene in a late step in the GA biosynthesis pathway (Sasaki et al 2002). ^4^ Many other short-culm cultivars were developed using an independent source of semidwarfism germplasms, such as the Japanese indigenous landrace ‘Jukkoku’ (Okada et al. 1967), or γ-ray-induced semidwarf cultivars such as ‘Reimei’ in Japan (Futsuhara et al. 1967) and ‘Calrose 76’ in the USA (Foster and Rutger 1978). Although dwarf varieties of rice have contributed to the dramatic improvement and stabilization of yields worldwide, the dwarf stature of varieties developed independently by using different native or mutant maternal lines happens to be controlled by a single dwarf gene, *sd1* (Sasaki et al. 2002; Kikuchi et al. 1997; Monna et al. 2002; Spielmeyer et al. 2002) The *sd1* gene confers the semidwarf phenotype with no detrimental effects on grain yield (Hedden 2003a; 2003b; Tomita and Ishii 2018), but as it is oligopoly by *sd1*.

Both *d35* from Tanginbouzu, which became the best rice breeding in Japan between 1955 and 1964, and *d18* from Kotake-tamanishiki were also kaurenoic acid oxidase- or 3-beta hydroxylase-defective in the same GA biosynthesis pathway (Itoh et al. 2004). On the other hand, *Rht*, which is related to the ‘Green Revolution’ of wheat, is a missing gene involved in GA’s signaling (Peng et al. 1999), and the Daikoku type d1 dwarf gene in rice is defective in the alpha subunit of the heterotrimeric G protein, affecting GA signal transduction (Ueguchi-Tanaka et al. 2000). Other dwarf genes such as *d11* (Tong et al. 2009)^17^, *sd37* (Zhang et al. 2014)^18^, whose function is not related to GA biosynthesis pathway, were certainly identified. However, their practical uses in breeding have not proceeded yet. A little genetic source of current semidwarf rice cultivars has a risk for environmental change. It is thus necessary to acquire a wider range of semidwarfing genes to cope with future environmental changes.

In order to find a new dwarf gene comparable to *sd1*, the first author conducted gene analyses focusing on Hokuriku 100, which is a mutant line with culms approx. 15 cm shorter than those of variety Koshihikari, and discovered a novel dwarf gene, *d60*, which brings about a good plant type with erect leaves by shortening culms by about 20%. Furthermore, and that it complements the game tic lethal gene, gal, to cause gamete lethality (Tomita 2012; Tomita and Tanisaka 2019). In other words, in the F1 hybrid (genotype *D60d60Galgal*) of Koshihikari (*D60D60galgal*) ×Hokuriku 100 (*d60d60GalGal*), male and female gametes having both gal and d60 become gamete lethal and the pollen and seed fertility decrease to 75%. As a result, F2 progeny shows a unique mode of inheritance that is segregated into a ratio of 6 fertile long-culm (4*D60D60*:2*D60d60GalGal*: 2 partially fertile long-culm (*D60d60Galgal*=F_1_ type):1 dwarf (*d60d60GalGal*) (Tomita 2012; Tomita and Tanisaka 2019). Furthermore, isogenic lines that were introduced with both *d60* and *sd1* genes into Koshihikari by backcrossing, namely, the d60sd1 line, became an additively extreme dwarf, suggesting that *d60* is functionally independent from *sd1* and not related to the GA1 biosynthesis pathway. The gamete sterilities caused by *gal* and *d60* in the hybrid between the original variety and its mutant were quite different from hybrid sterilities generally controlled by one-locus sporo-gametophytic allelic interaction at the single *S* locus (Kitamura 1962; Ikehashi 1985; Ikehashi and Araki 1986; Ikehashi 1991; Sano 1994), which identified from crosses between distantly related species with reproductive barriers. Therefore, it is intensely interested in the gene sequence and function of *d60*, which relates to both semidwarfness and gamete development.

## Materials and methods

### Chromosomal localization of *d60* by using morphological genetic marker

In order to determine the chromosomal location of the dwarf gene, *d60*, we conducted genetic linkage analyses of *d60* on the basis that the segregation ratios of the marker genes linked to *d60* do not fit the Mendelian ratio of 3:1. For the analyses, we developed F2 hybrids of 22 marker gene lines and Koshihikari d60 lines (Koshihikari*6//Koshihikari/Hokuriku 100), which were selected to cover all chromosomes as shown in Table 1. Taking into account the expectation that the segregation ratios for the marker genes linked to them in F2 differ from the Mendelian ratio of 3:1. In other words, when a recessive marker gene is fully linked to *D60*, the F_2_ segregation ratio of wild type to marker gene homozygous type will be 5:4. For each test cross combination between maker gene lines and Koshishikri d60 line, 300-400 F_2_ plants were grown at the Field Science Center. Seedlings were individually transplanted into a paddy field with density 22.2 seedlings/m^2^ (one seedling per 30 × 15 cm). The paddy field was fertilized by 4.0 kg of basal fertilizer containing nitrogen, phosphorus, and potassium (weight ratio, nitrogen:phosphorus:potassium = 2.6:3.2:2.6) at the rate with 4.3 g/m^2^ nitrogen, 5.3 g/m^2^ phosphorus, and 4.3 g/m^2^ potassium evenly across the field. Each F_2_ plants were phenotypically scored for each morphological marker gene and investigated for culm length and days to heading. The culm length between the ground surface and the panicle base of the main culm was measured for all plants. The time when the tip of the first panicle emerged from the flag leaf sheath was recorded as the heading time for all plants.

**Table 1.** Segregation ratio of morphological marker genes in F_2_ between the marker gene lines (*D60D60galgal*) and Koshihikari d60 line (*d60d60GalGal*)

| Marker genes<br>(chromosome) | Phenotypic segregation ratio<br>Wild type : marker gene homozygotes |  | X<sup>2</sup>(3:1) | Probability |
| --- | --- | --- | --- | --- |
| *d18* (1) | 317 | : 96 | 0.68 | 0.250<P<0.500 |
| *fs2* (1) | 318 | : 95 | 0.73 | 0.250<P<0.500 |
| *lax* (1) | 322 | : 104 | 0.08 | 0.750<P<0.900 |
| *Pn* (1) | 124 | : 304 | 3.60 | 0.050<P<0.100 |
| *d30* (2) | 195 | : 122 | 37.7 | P<0.005 |
| *gh2* (2) | 218 | : 100 | 7.05 | P<0.005 |
| *d1* (3) | 116 | : 35 | 0.27 | 0.500<P<0.750 |
| *bc2* (3) | 266 | : 66 | 4.64 | 0.010<P<0.025 |
| *lg* (4) | 222 | : 59 | 2.41 | 0.100<P<0.250 |
| *bgl* (5) | 116 | : 33 | 0.58 | 0.250<P<0.500 |
| *spl7* (5) | 111 | : 38 | 0.04 | 0.750<P<0.900 |
| *nl1* (5) | 317 | : 98 | 0.43 | 0.500<P<0.750 |
| *ri* (5) | 302 | : 113 | 1.10 | 0.250<P<0.500 |
| *d6* (7) | 199 | : 82 | 2.16 | 0.100<P<0.250 |
| *g* (7) | 195 | : 86 | 4.69 | 0.025<P<0.050 |
| *Rc* (7) | 63 | : 218 | 1.00 | 0.250<P<0.500 |
| *spl5* (10) | 98 | : 28 | 0.52 | 0.250<P<0.500 |
| *rfs* (11) | 100 | : 26 | 1.28 | 0.750<P<0.900 |
| *spl10* (11) | 189 | : 81 | 3.60 | 0.050<P<0.100 |
| *la* (11) | 274 | : 59 | 9.75 | P<0.005 |
| *sp* (11) | 259 | : 72 | 1.87 | 0.100<P<0.250 |
| *d28* (11) | 245 | : 84 | 0.05 | 0.750<P<0.900 |

### Molecular fine mapping of *d60*

A chromosome segment substitution line SL204 (*D60D60galgal*), which carries a 60.3-cM distal end segment of the short arm of the indica cultivar ‘Kasalath’ chromosome 2 to replace the corresponding d60-bearing segment in the background of the japonica cultivar ‘Koshihikari’, was crossed with a d60 Koshihikari line(*d60d60GalGal*). F_2_ plants (n=5122) were grown in the nursery cabinets, and short-culm plants (n=532) indicative of those homozygous for d60(*d60d60GalGal*) were selected by the round leaf shape of the young seedlings. The *d60* homozygous segregants were then transplanted individually at the Field Science Center. Each F_2_ plants were phenotypically scored and measured for culm length and days to heading. *d60* homozygous plants were confirmed again by the culm length and phenotypic characterization with erect leaves. The leaves were sampled individually from all F_2_ plants and powdered while being frozen in liquid nitrogen using a Precellys 24 high-throughput bead-mill homogenizer (Bertin Technologies, Montigny-le-Bretonneux, France), then the genomic DNA was extracted using the cetyl trimethylammonium bromide (CTAB) method. Forty-eight SSR markers on the short arm of chromosome 2 showing polymorphisms between Kasalath and Koshihikari were used to delimit the candidate region of *d60*. Using 20 ng each of rice genomic DNA as a template, 50 μL of a reaction solution containing 200 nmol/L forward and reverse primers (33 ng), 100 μmol/L dNTPs, 50 mmol/L KCl, 10 mmol/L Tris-HCl (pH 8.8), 1.5 mmol/L MgCl2, and 1 U TaKaRa LA Taq (Takara Bio Inc., Kyoto, Japan) was prepared. Using the Thermal Cycler CFX96 (BioRad Laboratories, Hercules, CA), the reaction solution was subjected to 35 cycles of denaturation at 94 °C for 30-s, annealing at 55 °C for 30-s, and primer extension at 72 °C for 1 min. The first denaturation at 94 °C and the last extension at 72 °C were set for 5 min. The SSR polymorphisms in the PCR products were analyzed by electrophoresis using a cartridge QIAxcel DNA Screening Kit (2400) in the QIAxcel electrophoresis apparatus (Qiagen, Hilden, Germany) at 5 kV for 10 minutes.

### Whole genome sequencing analysis

Whole genome sequencing was conducted on Koshihkari d60(BC_8_F_2_), by eight backcrosses into the genetic background of Koshihikari. The leaves were powdered using a mortar and pestle and frozen in liquid nitrogen. Genomic DNA was extracted from each cultivar using the cetyltrimethylammonium bromide method. Genomic DNA was fragmented and simultaneously tagged so that the peak size of the fragments was approximately 500 bp using the Nextera® transposome (Illumina Inc., San Diego, CA). After purification of the transposome using DNA Clean & ConcentratorTM-5 (Zymo Research, Irvine, CA), adaptor sequences, including the sequencing primers, for fixation on the flow cell were synthesized at both ends of each fragment using polymerase chain reaction. The DNA fragments were then subjected to size selection using AMPure XP magnetic beads (Beckman Coulter, Brea, CA). Finally, qualitative checks were performed using a Fragment Analyzer™ (Advanced Analytical Technologies, Heidelberg, Germany) and quantitative measurements using Qubit® 2.0 Fluorometer (Life Technologies; Thermo Fisher Scientific, Inc., Waltham, MA) to prepare a DNA library for next-generation sequencing. Sequencing was conducted in paired-end 2 × 100 bp on a HiSeq X next-generation sequencer, according to the manufacturer’s protocol (Illumina Inc., San Diego, CA). Illumina reads were trimmed using Trimmomatic (version 0.39) (Bolger et al. 2014) (Figure 1). Sequencing adapters and sequences with low quality scores on the 3′ ends (Phred score [Q], < 20) were trimmed. The raw Illumina whole genome sequence reads were quality checked by performing quality control using FastQC (version 0.11.9; Babraham Institute). Mapping of reads from Koshihikari d60 to the Koshishikri genome as a reference was conducted using Burrows-Wheeler Aligner software (version bwa-0.7.17.tar.bz2) (Li and Durbin 2014). Duplicated reads were removed using Picard (version 2.25.5) (http://broadinstitute.github.io/picard) and secondary aligned reads were removed using SAMtools (version 1.10.2) (Li et al. 2009). To identify genetic variations among strains, single nucleotide variant detection (variant calling) and single nucleotide variant matrix generation were performed using GATK (version 4.1.7.0) (McKenna et al.).

**Fig. 1.**
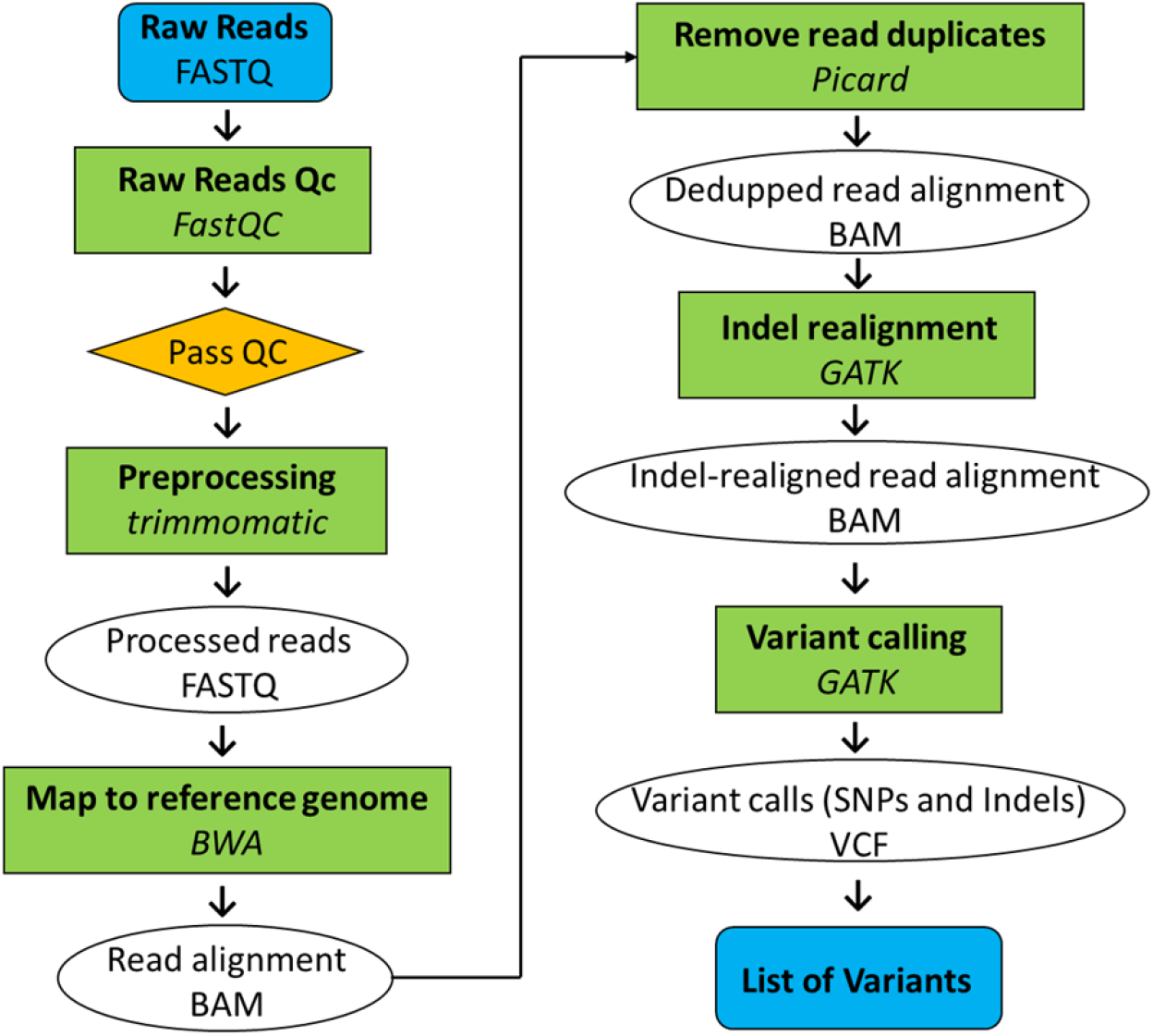
Pipeline of whole genome resequencing analysis.

### RT-qPCR analysis

Leaves were sampled from Koshishikari and Koshihkari d60 (BC7F3) before anthesis. Total RNA was extracted from the leaves using the Acid Phenol Guanidium Chloroform method (Chomczynski and Sacchi 2006) and was then treated with 10 U of DNase I (Takara Bio Inc., Kyoto, Japan). poly(A)+ RNA was separated and purified from total RNA using an oligo(dT) cellulose column Poly(A) Quick mRNA Isolation Kit (Stratagene, La Jolla, CA). Single-stranded cDNAs were synthesized by reverse transcription reaction using 0.25 U AMV reverse transcriptase (Life Sciences Advanced Technologies, Saint Petersburg, FL) for 30 min at 50°C by using 200 ng of poly(A)+RNA as the template of 2.5 μM oligo(dT) primers in 20 μL solution (1 mM dNTP, 5 mM MgCl_2_. 10 mmol/L Tris-HCl (pH 8.3), 50 mmol/L KCl). For RT-qPCR, we designed a FAM-labeled probe from three candidate genes for *d60* located around 10.3 Mb that may be involved in height growth, namely, an entocopalyl disphosphate synthase-like gene that acts upstream of gibberellin biosynthesis and xyloglucan endotransglucosylase-like genes involved in the structure of the cell wall. 18srRNA was used as an endogenous control. The reaction mixture contained 25 µg of total RNA of Koshishikari or Koshihkari d60 (BC_7_F_3_), 10 µL of Prime Time Gene Expression Master Mix (2x), and 1 µL of Prime Time qPCR Assay (20x) in a volume of 20 μL. 45 cycles of RT-qPCR comprising denaturation at 95 °C for 15 s and annealing/extension at 60 °C for 45 s were conducted and the fluorescence was detected with the thermal cycler CFX96.

## Results

### Distorted segregation of genetic markers on chromosome 2 in 5:4 from Mendelian ratio 3:1

We performed genetic linkage analyses based on the expectation that the segregation ratios of the marker genes linked to *d60* do not fit the Mendelian ratio of 3:1 (Fig. 2). Namely, 22 morphological marker gene lines were crossed with the Koshihikari d60 line (Koshihiakri *6//Koshihikari / Hokuriku No. 100) (Table 1). Among the F_2_ with 22 marker gene lineages that have *d30* on chromosome 2 (*D60D60galgal*) and Koshihikari d60 line (*d60d60GalGal*), the segregation ratio of wild type to *d30* homozygotes at the *d30* locus was 195:122 (Table 1), which deviated significantly from 3:1, but it was closer to the theoretical ratio of 5:4 (χ^2^=4.56, 0.025<P<0.05) for the cases where *d30* is fully linked to *D60*. Furthermore, also at *gh2* locus the ratio of wild type to *gh2* homozygote was 218:100 (Table 1), which deviated significantly from 3:1, indicating a linkage between *d60* and *gh2* on chromosome 2.

**Fig. 2.**
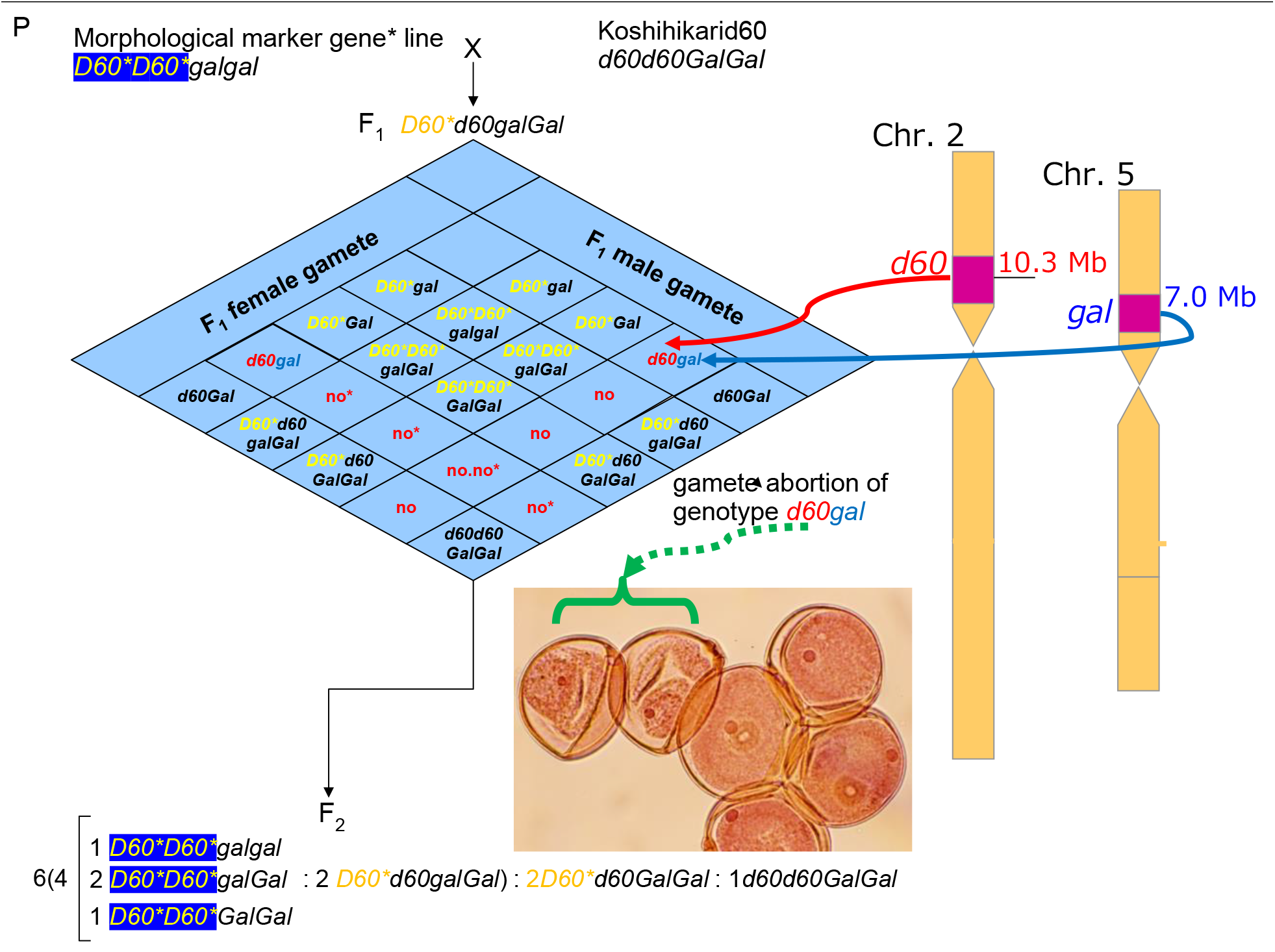
Identification of chromosomal location of *d60* by distorted segregation of morphological marker genes linked to *D60*. Gamete abortion caused by *d60* on chromosome 2 and *gal* on chromosome 5. Pollen with genotype *d60gal* began to degrade at the binucleate stage and degraded pollen lost vegetative nuclei, but a second pollen mitosis was occurred raising two generative nuclei (Tomita and Tanisaka 2019). If the recessive morphological gene is tightly linked to *D60*, the segregation ratio of wild type : recessive homozygotes is deviated to 5:4 from the Mendelian 3:1 ratio.

### Molecular fine mapping of *d60* on chromosome 2

A fine mapping was performed by using *d60* homozygous (*d60d60GalGal*) plants segregated in F_2_ between line SL204 (*D60D60galgal*), in which chromosome 2 of Koshihikari was substituted by the indica variety Kasalath, and the Koshihikari d60 line (*d60d60GalGal*). we transplanted 532 segregants of the *d60* homozygote to the Field Science Center. We used SSR markers on chromosome 2 that show DNA polymorphisms between Koshihikari and Kasarash to locate the *d60* DNA sequence. As a result, 8 markers RM12918, RM452, RM12938, RM12949, RM1358, RM12970, Os02ssr, and RM13002, which lie at 9.5, 9.6, 9.9, 10.1, 10.2, 10.3, 10.5, and 10.7 Mb from the distal end of the short arm of chromosome 2, showed a recombination value of 5.7, 5.1, 3.9, 2.1, 3.2, 0.0, 0.7, and 4.5, respectively (Fig. 3). The results showed a close linkage of *d60* with RM12970 on chromosome 2 with a recombination value of 0.0, suggesting the location of the *d60* locus is in the region 10.3 Mb away from the distal end of the short arm of chromosome 2.

**Fig. 3.**
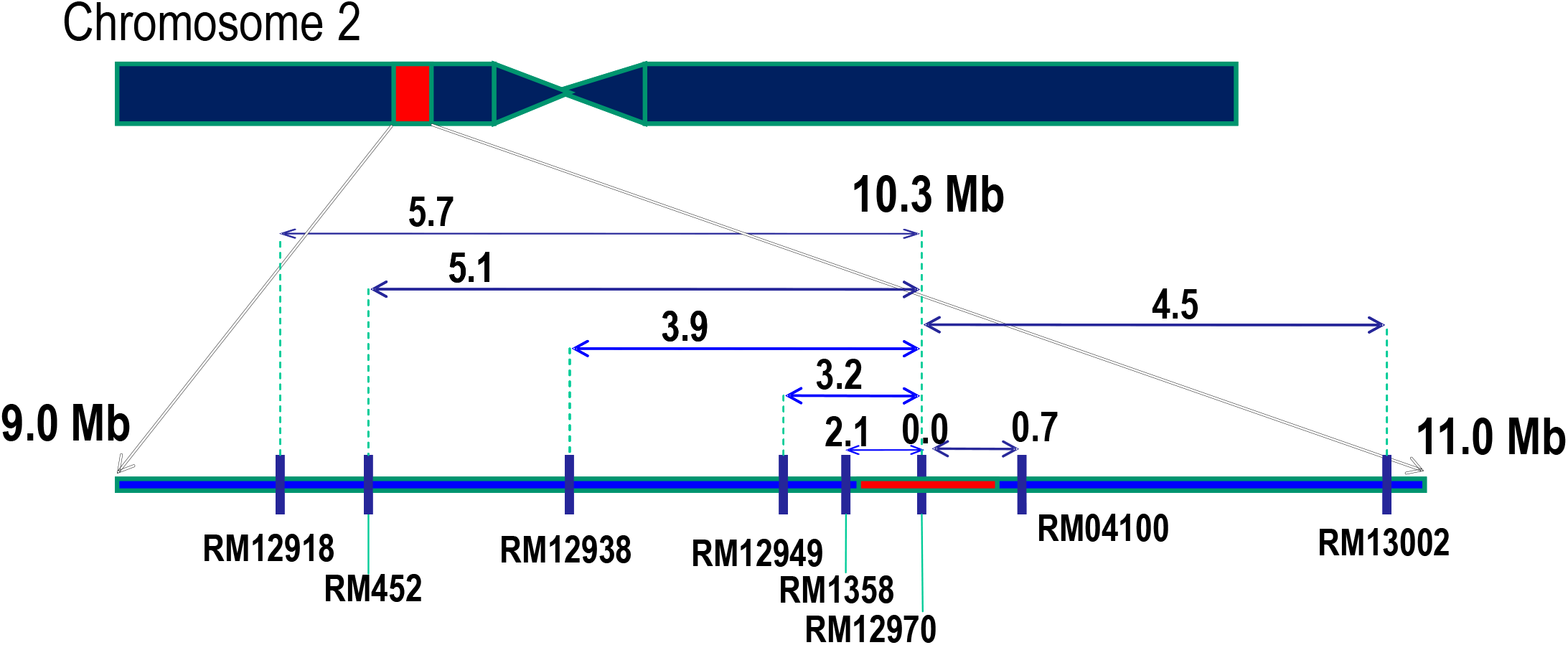
Molecular fine mapping of semidwarfing gene *d60*. A close linkage of *d60* with RM12970 with a recombination value 0.0 was detected at 10.3 Mb away from the distal end of the short arm of chromosome 2.

### Whole genome sequencing of Koshihikari d60

The number of reads decoded by the next-generation sequencer was 126,884,326 for Koshihikari d60 (BC_8_F_2_). The gained reads of Koshishikri d60 were mapped to the consensus sequence of Koshihikari as the reference (No. of mapped reads 126,733,901; % of Mapped reads 99.88%), and the mean coverage were 22.42. After removing the secondary alignment and duplicate reads, the unique reads were 122,934,478. A total of 43,861 SNPs [homozygous 2044, heterozygous 41817] were detected. The number of SNPs were less than 10 per 0.1 Mb through the genome (Fig. 4). The results indicated that a large portion of the rice 12 chromosomes were substituted to the genome of Koshihikari after continuous backcross targeting these two genes. By molecular fine mapping, the candidate location of *d60* was narrowed around 10.3 Mb away from the distal end of the short arm of chromosome 2. However, whole genome sequencing of Koshihikari d60 revealed no SNPs and Indels around 10.3 Mb region, which were tightly linked to *d60*. No. of SNPs

**Fig. 4.**
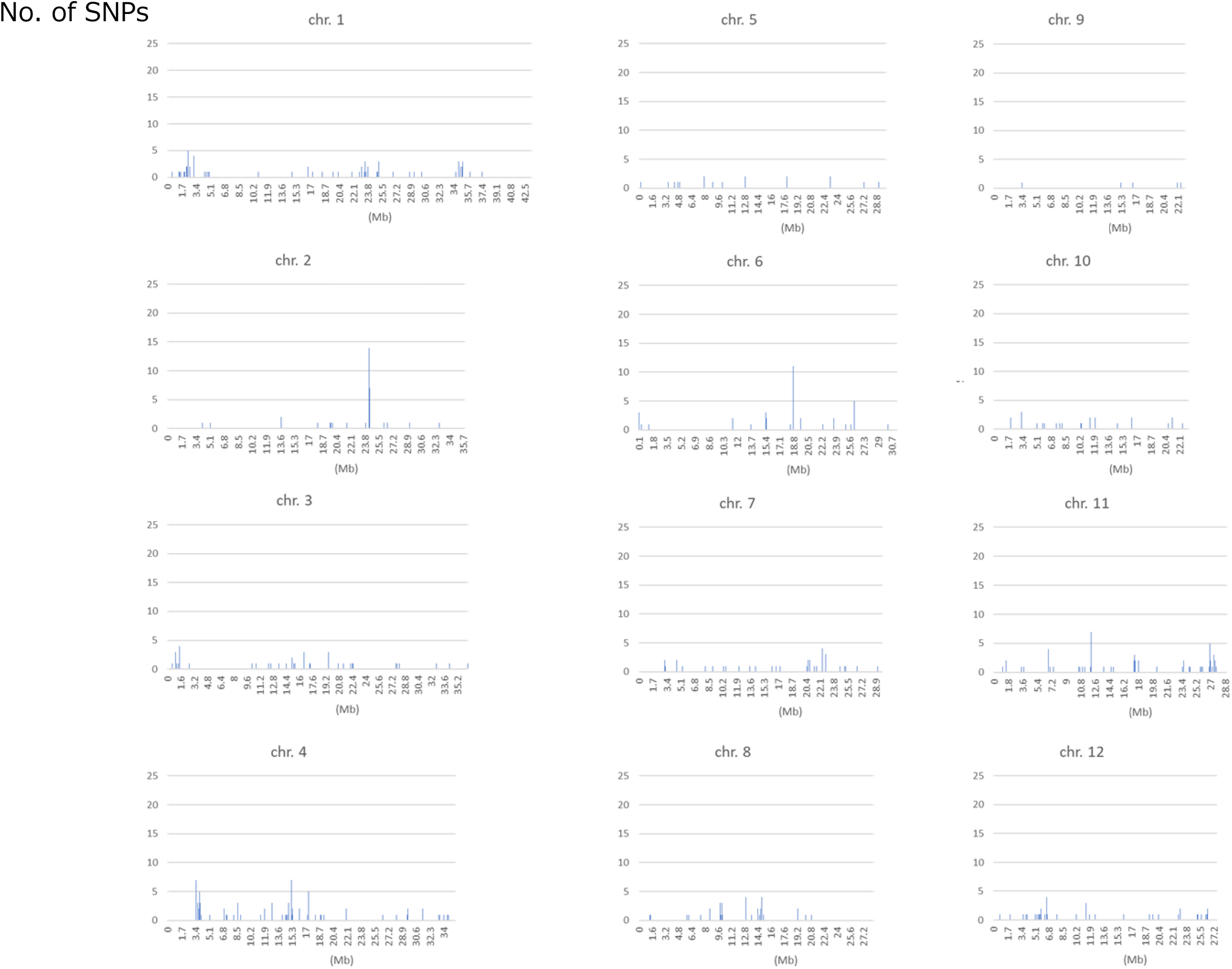
Frequency distribution of SNPs per 0.1 Mb detected by whole genome sequencing of Koshihikari d60 (B_7_F_6_) based on SNV calling against the reference genome of Koshihikari. There were at most less than 5 SNPs per 0.1 Mb over all genomes.

### RT-qPCR analysis of xyloglucan endotransglucosylase gene

Fine mapping using DNA markers localized the short culm gene *d60* to approximately 10.3 Mb from the end of the short arm of chromosome 2, but no mutations were detected in the genomic DNA in that region. Therefore, we performed RT-qPCR to analyze whether there were any differences in the expression levels of three candidate genes for *d60* located around 10.3 Mb that may be involved in height growth, namely, an entocopalyl disphosphate synthase-like gene that acts upstream of gibberellin biosynthesis and xyloglucan endotransglucosylase-like genes involved in the structure of the cell wall. As a result, both in Koshihikari d60 and Koshihikari d60Hd16, of the three candidate genes, there was no difference in expression between the two xyloglucan glucosyltransferase-like genes compared to Koshihikari (Fig. 5B, C), but only the transcription of the ent-copalyl diphosphate synthase-like gene was clearly lower than that of Koshihikari (Fig. 5A). The reduced expression of this gene namely by *d60* allele was thought to cause short culms.

**Fig. 5.**
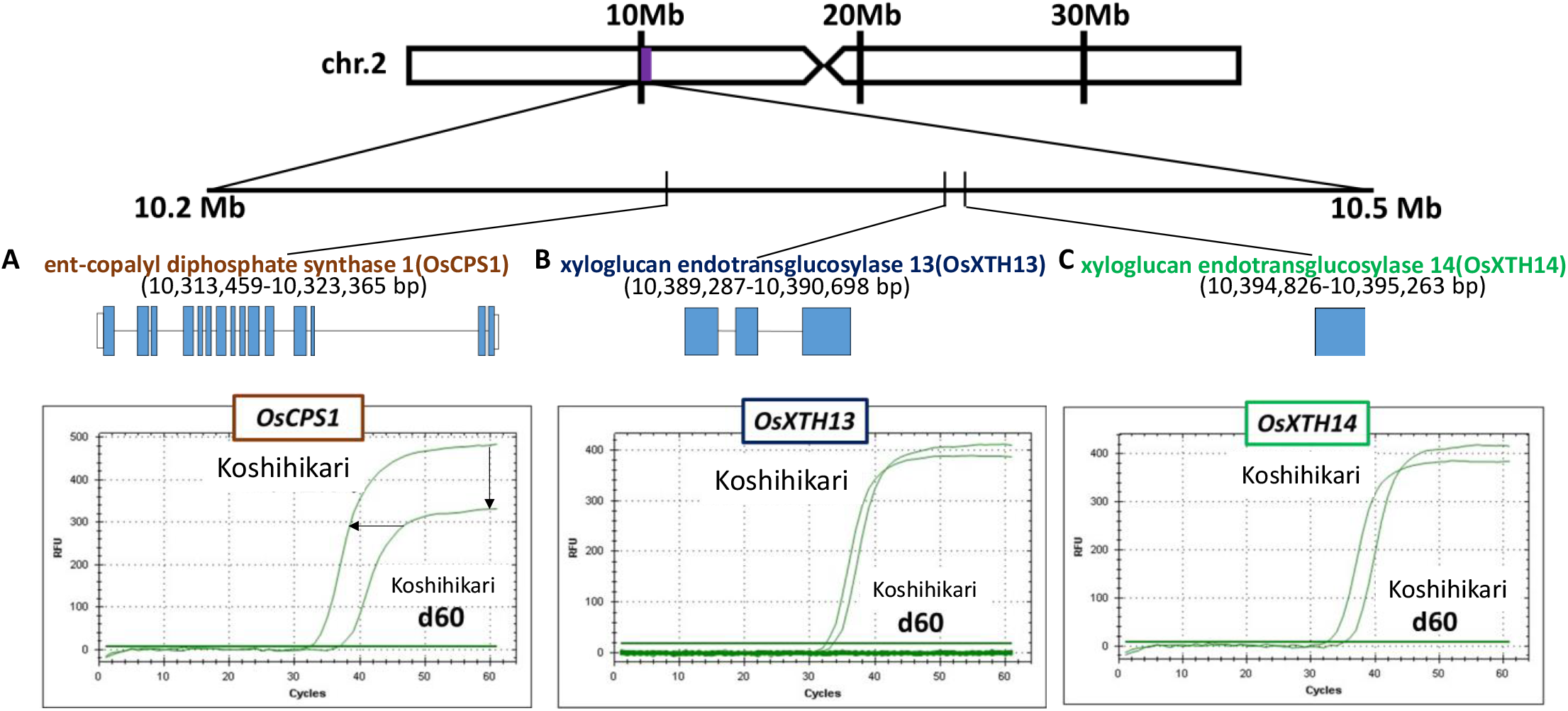
RT-qPCR for three candidate genes for *d60* located around 10.3 Mb that may be involved in height growth. **A** RT-qPCR results of ent-copalyl diphosphate synthase-like gene indicated that the rising curve of the fluorescence signals in Koshihikari d60 was apparently reduced than that in Koshihikari. B, C In Koshihikari d60, there was no difference in expression between the two xyloglucan glucosyltransferase-like genes compared to Koshihikari.

## Discussion

The threat of strong typhoons is increasing, probably as a result of global warming (Knutson et al. 2019). This is a serious problem in rice production, because strong winds cause stem lodging and consequent yield losses and deterioration in crop quality (Dahiya et al. 2018). The semidwarf character is one of the essential agronomic traits in crop breeding. Extensive damage from lodging of rice due to frequent typhoons has become a social problem in recent years, and developing new varieties of typhoon-resistant rice by introducing dwarf genes is an urgent task. Hence, there is a pressing need to develop new short-culm rice cultivars resistant to strong winds (Hirano et al. 2017). Breeding of dwarf varieties brought about an improvement in lodging resistance and epoch-making yield quantity, as represented by Green Revolution. The breeding program that has made the greatest contribution in the history of mankind is the ‘Green Revolution,’ in which the production of grain was dramatically increased in the 1960s with the development of dwarf varieties of rice. These dwarfs are brought by only a single kind of semidwarf gene *sd1*. The *sd1* allele is located on the long arm of chromosome 1 (Cho et al. 1994a, b; Maeda et al. 1997), which confers the semidwarf phenotype without detrimental effects on grain yield (Hedden 2003a, b). However, in consideration of the purpose of attempting to maintain/expand the genetic diversity of varieties, so we should not rely on the *sd1* gene only. Namely, the use of sd1 and other genes defective in the same GA biosynthesis pathway are after all merely controlling the amount of GA1, which is a final product of active gibberellin, in the stem and leaf.

In order to look for a novel dwarf gene that can take the place of the green revolution gene sd1, the first author focused on radiation-induced short stem mutant Hokuriku 100, which is a short stem mutant line with culms approx. 15 cm shorter than those of the original variety Koshihikari. Despite the promising phenotype of Hokuriku 100, it has been left aside from practical breeding, because of the unclear segregation of semidwarfness. The first author conducted gene analyses on Hokuriku 100 and discovered a novel dwarf gene, *d60*, which confers an excellent plant type with shortened culms by approx. 20% and erect leaves (Tomita 2012; Tomita and Tanisaka 2019). When the first author crossed this dwarf mutant with Koshihikari, instead of exhibiting the conventional Mendel’s Law, that is, a 3:1 ratio for long:short culm lengths, the F_2_ generation of the dwarf variety and Koshihikari had a 8:1 ratio and produced only a very small number of small-size rice plants (Tomita 2012; Tomita and Tanisaka 2019). Further investigation found that the F_2_ population of tall rice contained a certain amount of partial sterile plants that did not bear seeds (Tomita 2012; Tomita and Tanisaka 2019). Finally, it is concluded that in the F_1_ hybrid (genotype *D60d60Galgal*) of Koshihikari (*D60D60galgal*)×Hokuriku 100 (*d60d60GalGal*), male and female gametes having both gal and d60 become gametic lethal and the pollen and seed fertility decrease to 75%. As a result, F_2_ progeny shows a unique mode of inheritance that is segregated into a ratio of 6 fertile long culm (4*D60D60*: 2*D60d60GalGal*):2 partially fertile long culm (*D60d60Galgal*=F_1_ type):1 dwarf (*d60d60GalGal*) (Tomita 2012; Tomita and Tanisaka 2019).

In this study, according to the expectation that the segregation of marker genes linked with *d60* deviates from the Mendelian ratio of 3:1, a linkage analysis was conducted between 22 marker gene lines and *d60*. Among the marker genes, *d30* and *gh2* on chromosome 2 showed an excessive segregation, which was close to the segregation ratio of 5:4 in the case of complete linkage to *D60*, and *d60* was localized on the short arm of chromosome 2. Furthermore, the double dwarfness by combining *d60* and *d30* was obtained in F_3_ via recombination breaking repulsion-phase linkage on rice chromosome 2 (Tomita and Tanaka 2019). *d60* was also linked with a large grain gene *GW2* on chromosome 2 (Tomita et al. 2019). Next, the Koshihikari d60 line was crossed with a chromosome segment substitution line that carried a segment of chromosome 2 derived from Indica cultivar ‘Kasalath’ in the background of the *japonica* cultivar ‘Koshihikari’. Short-culm homozygous (*d60d60GalGal*) plants in the resulting F_2_ progenies were examined for genetic linkage by using SSR markers located on the short arms of chromosomes 2, thereby achieving a fine mapping resolution of *d60*. As a result, the locus of *d60* was narrowed down to 10.3 Mb region from the end of the short arm of chromosome 2 (recombination value 0.0). Analysis of the DNA sequences in the vicinity of the *d60* gene revealed the presence of ent-copalyl diphosphate synthase-like gene and RT-qPCR analysis showed its reduced expression. These mutations on the independent chromosomes happened by chance via gamma-ray irradiation.

The complementary gamete sterility caused by *gal* and *d60* is similar to that proposed in hybrid sterilities between subspecies of rice, which were explained by the two-gene model. Namely, the duplicate *S* gene loci causes hybrid male sterility when the F_1_ male gametes receive both recessive *S* genes on each duplicate locus (Oka 1953, 1974). For example, if japonica parent A and indica parent B have genotypes *s1/s1+2/+2* and *+1/+1s2/s2*, respectively, in which at least one + gene is necessary for male gamete development, therefore 25% of the F_1_ will be pollen sterile. Because *gal* and *d60* cause both sex sterilities in japonica hybrids, these genes were different from the duplicate *S* gene model, only causing male sterility in intersubspecies hybrids. Pollen with genotype *d60gal* began to degrade at the binucleate stage and degraded pollen lost vegetative nuclei, however, a second pollen mitosis occurred raising two generative nuclei (Tomita and Tanisaka 2019).

Hokuriku 100 has been regarded as a promising semidwarf strain with an excellent plant type with erect leaves (Tabuchi et al 2000). However, Hokuriku100 has never been used for practical breeding except for the cultivar Minihikari (Tomita 2012), because the inheritance mode of the semidwarf gene *d60* is complicated (Tomita 2012; Tomita and Tanisaka 2019) and the responsible DNA sequence was not identified. Then, the first author developed an isogenic line which had both *d60* and *sd1* by introducing both genes into Koshihikari by backcrossing and that the Koshihikari d60sd1 line became an additively extreme dwarf, demonstrating that *d60* and *sd1* are functionally independent (Tomita 2012) and not related to the GA1 biosynthesis pathway. In this study, the responsible DNA for *d60* was inferred as an ent-copalyl diphosphate synthase-like gene. In addition, regarding to the yield performance, *d60* can be as effective as *sd1* (Tomita and Ishimoto 2019). Therefore, *d60* would be expected to become one of the alternatives to *sd1* among the limited semidwarf gene pool.

In Japan, Koshihikari is a major rice cultivar and has approximately 40% of rice acreage, and even 80% in certain regions (Rice Stable Supply Organization). However, there is a large risk if adverse events should occur. Koshihikari has shown a reduction in yield and loss of quality due to lodging and immature chalky grains, which have been caused by recent climate change due to global warming. Therefore, there is an urgent need to develop a semidwarf Koshihikari variety that is resistant to lodging. The author developed a dwarf Koshihikari-type cultivar, ‘Hikarishinseiki’ (Tomita 2009; MAFF 2004) (rice cultivar No. 12273, Ministry of Agriculture, Forestry and Fisheries of Japan) The dwarf cultivar was developed by intercrossing the F_4_ of Kanto No. 79 (early-heading mutant line derived from Koshihikari) and Jukkoku (a cultivar with a semi-dwarf gene, *sd1*) as the maternal dwarf parent using with the Koshihikari-type heading time, and backcrossed it with Koshihikari 8 times. Having over 99.8% background of the Koshihikari genome, except for *sd1*, Hikarishinseiki would be the first cultivar to be registered as a Koshihikari-type dwarf with *sd1* in Japan (Tomita 2009). Through this study, we identified a novel semidwarf gene *d60* on chromosome 2, which is genetically and functionally independent from *sd1. d60* confers a good semidwarf phenotype without detrimental effects on grain yield (Tomita & Ishimoto 2019). Furthermore, a semidwarf and large grain isogenic Koshihikari was developed by integrated with *d60* and *GW2* (Tomita et al. 2019). Therefore, *d60* is a useful gene source to diversify semidwarf rice varieties. We already established the diagnosis of Jukkoku_*sd1* by using *Pma*CI digestion of PCR products on the SNP (Naito and Tomita 2013; Tomita and Ishii 2018). In this study, we identified DNA markers, which are closely linked to *d60* and higher efficiency of selection, and provided promising semidwarf breeding by using both markers selecting methods of *sd1* and *d60*.

## Conclusions

Rice semidwarf gene *d60* was inherited together with a pleiotropic effect on gamete abortion. Namely, F_2_ progenies of Koshihikari (*D60D60galgal*, long stem) × Hokuriku 100 (*d60d60GalGal*, short stem) segregated in the ratio of 1 semidwarf (1 *d60d60GalGal*):2 tall and quarter sterile (2 *D60d60Galgal*):6 tall (2 *D60d60GalGal*:1 *D60D60GalGal*:2 *D60D60Galgal*:1 *D60D60galgal*). *d60* was analyzed by linkage with phenotypic marker genes, whose segregations did not fit Mendelian ratio. F_2_ of Koshihikari d60 line (*d60d60GalGal)* × lines carrying *d30* or *gh2* on chromosome 2, the segregation ratios of these alleles were deviated from 1:3 but well fitted to 5:4 when *d30* and *gh2* were closely linked to *D60*. Furthermore, molecular fine mapping of *d60* by using DNA polymorphisms in F_2_ of Koshihikari d60 line × line carrying the indica chromosome 2 segment in Koshihikari showed that *d60* was tightly linked with RM12970 by a recombination value 0.0 at the region 10.3 Mb of chromosome 2. However, whole genome sequencing of Koshihikari d60 revealed no SNPs and Indels around the 10.3 Mb region. RT-qPCR for ent-copalyl diphosphate synthase-like gene on 10.3 Mb indicated its transcription was reduced compared to that of Koshihikari.

## Acknowledgements

This work is founded on Adaptable and Seamless Technology Transfer Program (A-STEP) through Target-driven R&D (high-risk challenge type) by Japan Science and Technology Agency (JST) to Motonori Tomita, whose project ID14529973 was entitled “NGS genome-wide analysis-based development of rice cultivars with super high-yield, large-grains, and early/late flowering suitable for the globalized world and global warming”, since 2014 to 2018, and 2021 to 2022.

